# Analysis of Confounding Factors in Reactive Cysteine Profiling Reveals Enhanced Chromatin-Protein Association via CDK7 Inhibition by THZ1

**DOI:** 10.64898/2026.05.05.721470

**Authors:** Ka Yang, Shaoxian Li, Bohui Li, Daniel M. Richards, Kevin Dong, Uthpala Seneviratne, Wankyu Lee, Anthony Iannetta, Hua Xu, Steven P. Gygi, Qing Yu

## Abstract

Recent advances in activity-based proteome profiling (ABPP) have enabled global mapping of cysteine ligandability, uncovering novel biological insights and opportunities for identifying disease vulnerabilities. While both live cell-based and native lysate-based ABPP have been applied, how cysteine ligandability differs between these systems and what factors influence these measurements remain unclear. Building on our previous development of a high-throughput TMT-ABPP workflow for native lysates, here we adapt the protocol for live cells and systematically compare cysteine ligandability across both platforms. Our analysis reveals three major contributors to the discrepancies: in-cellulo cysteine accessibility, protein abundance changes, and protein relocalization. Notably, we highlight that CDK7 inhibitor THZ1 induces substantial protein relocalization and promotes chromatin binding. Together, these results provide a practical framework for ABPP experimental design and data interpretation, supporting more accurate application of ABPP in functional proteomics and drug discovery.

## Introduction

Proteins are essential functional molecules in an organism, carrying out almost all biological processes. Competitive activity-based protein profiling (ABPP) has emerged as a transformative technology for functional proteomics, enabling the direct assessment of protein activity using residue-directed covalent probes^1–4^. Originally developed to assess functional states of specific subsets of enzymes, such as serine hydrolases^5,6^ and cysteine proteases^7,8^, ABPP has rapidly evolved far beyond interrogating enzymatic residues into a powerful tool to globally assess many additional types of functionality and protein-ligand interactions for biomedical research and drug discovery^2,9–13^.

Among all residues, cysteine has gained growing traction. Its intrinsic nucleophilicity, sensitivity to redox and metabolic cues, and involvement in catalysis, metal coordination, and allosteric regulation make it an ideal hotspot for probing protein functional states^7,14,15^. The recent successful cases of covalent drug discovery has further fueled interest in cysteine profiling, notable examples including Sotorasib and Adagrasib which therapeutically target the once “undruggable” KRAS^G12C^ mutant in various types of cancer^2,7,9,16^. Consequently, cysteine-focused ABPP has become a powerful technology for mapping functional and druggable residues proteome-wide as well as discovery of electrophilic ligands.

Liquid chromatography-mass spectrometry (LC-MS)-based ABPP represents a major advance beyond gel-based formats by enabling proteome-scale quantification of cysteine reactivity and ligand engagement^17–19^. ABPP has been applied in small-scale drug screening^20,21^. However, the current throughput is incompatible with the immerse chemical space. To improve throughput and quantitative accuracy, tandem mass tag (TMT)-based sample multiplexing has been incorporated into ABPP using a desthiobiotin iodoacetamide (DBIA) probe, enabling up to 18 samples to be analyzed simultaneously with high quantitative accuracy and consistency^19,22–25^. We recently developed 96 well plate-based TMT-ABPP, enabling compound profiling using minimal proteome input (∼10 µg lysate per sample) at scale^19,23,26^. This miniaturized and high-throughput approach now makes it feasible to screen electrophile libraries while maintaining quantitative depth and proteome coverage, an essential step toward bridging functional proteomics with early-stage drug discovery.

ABPP has been traditionally performed using native cell lysate, in part because lysate-based workflows reduce sources of biological variability and provide robust, reproducible measurements of cysteine reactivity and ligandability^7,9,27^. However, the concern that lysing cells leads to the loss of cellular context, including membrane-bound organelles, protein-protein interactions, and protein modification state (e.g. phosphorylation), has motivated recent research using live cells^22,26,28,29^. While both lysate-based and live-cell approaches have yielded important biological discoveries, substantial discrepancies between the two systems have frequently been observed. The major drivers of these inconsistencies have remained unclear and thereby underestimated, posing challenges for experimental design, data interpretation, and reproducibility in the field^30^.

Here, we extend our TMT-ABPP platform to live-cell applications. In addition to providing detailed protocols optimized for both lysate-based and live-cell formats, we systematically investigate potential sources of discrepancy in cysteine ligandability across these systems. We show that three major factors, including 1) predicted cysteine accessibility, 2) rapid protein abundance change triggered by compound treatment, and 3) active protein relocalization, contribute to the observed inconsistency. We further highlight that THZ1, a CDK7 inhibitor, triggers substantial proteome remodeling by promoting the association of diverse proteins with chromatin. Together, these findings underscore the importance of accounting for cellular context, dynamic proteome responses, and subcellular proteome redistribution when designing ABPP experiments and interpreting results. Consideration of these factors will be essential for improving consistency, reproducibility, and biological insight across future functional proteomics studies.

## Results and Discussion

Competitive TMT-ABPP enables systematic and quantitative assessment of cysteine ligandability in the human proteome and can be applied either in live cells or native cell lysates^22,26–28^. Despite successes using either system, debates remain regarding reproducibility and biological relevance. Native lysates offer major practical advantages: they are straightforward to generate in large quantities, minimize biological variability, and are well suited for high-throughput screening. However, native lysis of cells, though mild, leads to the inevitable loss of cellular environments and membrane-bounded compartments, critical to proper protein folding and protein-protein interactions. On the other hand, live-cell ABPP, though preserving the intact cellular environment, is inherently more variable and susceptible to secondary effects induced by compound treatment that may confound the interpretation of cysteine ligandability. These factors complicate direct comparisons between the two systems and pose challenges for interpreting cysteine reactivity and ligandability.

We recently developed a high-throughput TMT-ABPP platform that enables the screening of hundreds of electrophilic compounds in 96-well plates using only 10–20 □g of native proteome lysate per well^19,23^. Here we expand it for cysteine profiling in live cells (**Fig. 1A**). Both systems offer comparable throughput, with live-cell profiling requiring only ∼30 min of additional time for cell lysis following compound incubation, whereas lysate-based profiling can use bulk native lysate prepared in advance. To investigate the causes of the discrepancies of ligandability observed between live-cell and lysate-based ABPP measurements, we systematically evaluated factors that influence the competition ratio (CR), using matched workflows across both systems. For clarity, we refer to CL (Cell-Lysate) conditions as compound treatment performed in living cells followed by DBIA incubation in lysate (native and soluble proteome)^22^, and to LL (Lysate-Lysate) conditions as both compound and DBIA incubation performed in native lysate^23^. We profiled cysteine engagement using 6 compounds with broad and diverse engagement patterns bearing two common covalent warheads (chloroacetamide and acrylamide)^9,23^, generating two TMT18 plexes for either system (**Fig. 1B and Fig. S1A**). KB02, KB03, and KB05 are widely used, highly reactive scout compounds^9,22,23^, while THZ1 represents a well-studied, selective CDK7 inhibitor^31^, and AC19 and CL4 represent poorly characterized compounds^22,23^. Both systems measured comparable number of reactive cysteines, confirming the successful adaptation of the TMT-ABPP platform in live cells (**Fig. 1C**). Measurements were precise, with median coefficients of variation (CVs) across replicates below 8% in both systems. As expected, CL exhibited slightly higher CVs than LL by ∼1% (**Fig. S1B**). The majority (70%) of measured cysteines were shared between the CL and LL datasets (**Fig. 1D**), providing a controlled basis for comparing ligandability differences across the two systems. Consistent with this overlap, cysteine accessibility, estimated by the prediction-aware part-sphere exposure (pPSE) score^32,33^, was highly comparable between the two systems (**Fig. 1E**).

**Figure 1.**
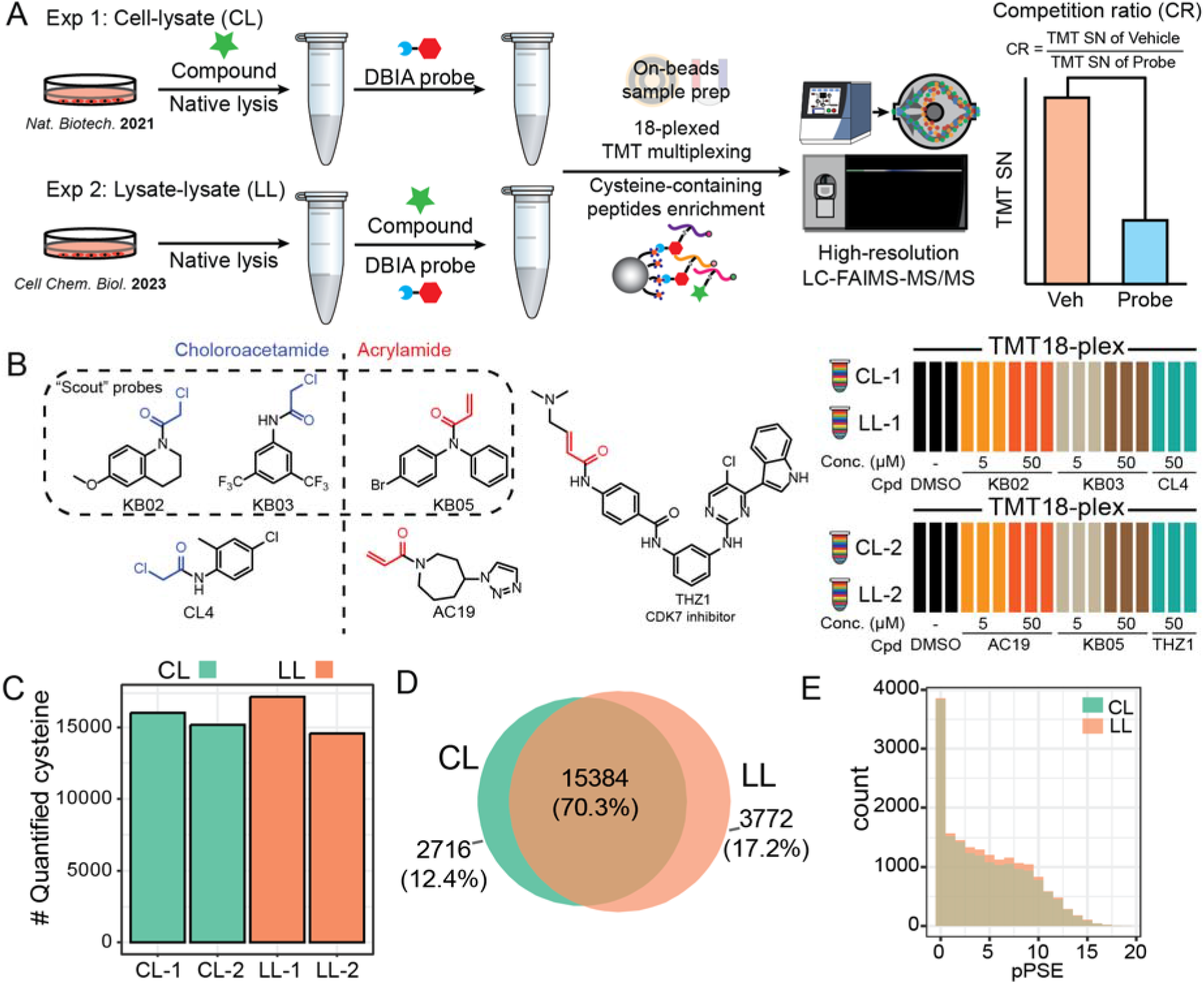
Two common reactive cysteine profiling strategies are evaluated. **A)** Workflow to compare two ABPP strategies. Cell-Lysate (CL) stands for in-cell probe treatment compound treatment followed by lysis and incubation with DBIA probe. Lysate-Lysate (LL) stands for in-lysate compound treatment followed by incubation with DBIA probe. Competition ratio (CR) calculated from TMT signal-to-noise (S/N) values was used to evaluate ligandability. **B)** Covalent probes used in this work and TMT experiment design. **C)** Quantified numbers of unique reactive cysteines in each experiment. **D)** Overlap of quantified cysteine by two strategies. **E)** pPSE accessibility score distribution of DBIA-labeled reactive cysteine residues.

However, despite significant overall correlation between CR^CL^ and CR^LL^ (R = 0.36), direct comparison of the two datasets reveals clear discrepancies (**Fig. 2A**). Among ∼1,700 ligandable cysteines (CR ≥□2 by any compound), only 148 (9%) are common to both CL and LL, such as TADA3 C255, C5orf51 C179 and INTS13 C349, and a large fraction of ligandable sites are system-specific (**Fig. 2B, 2C, and S2A**). It is clear to us that the LL workflow generated more ligandable cysteine measurements (**Fig. 2C and S2A**), suggesting that lysate conditions expose additional cysteine that are inaccessible in intact cells. At the protein level, ∼17% of 1,027 ligandable proteins were consistently liganded across both systems, representing a higher overlap than observed at the cysteine level (9%) (**Fig. 2C**).

**Figure 2.**
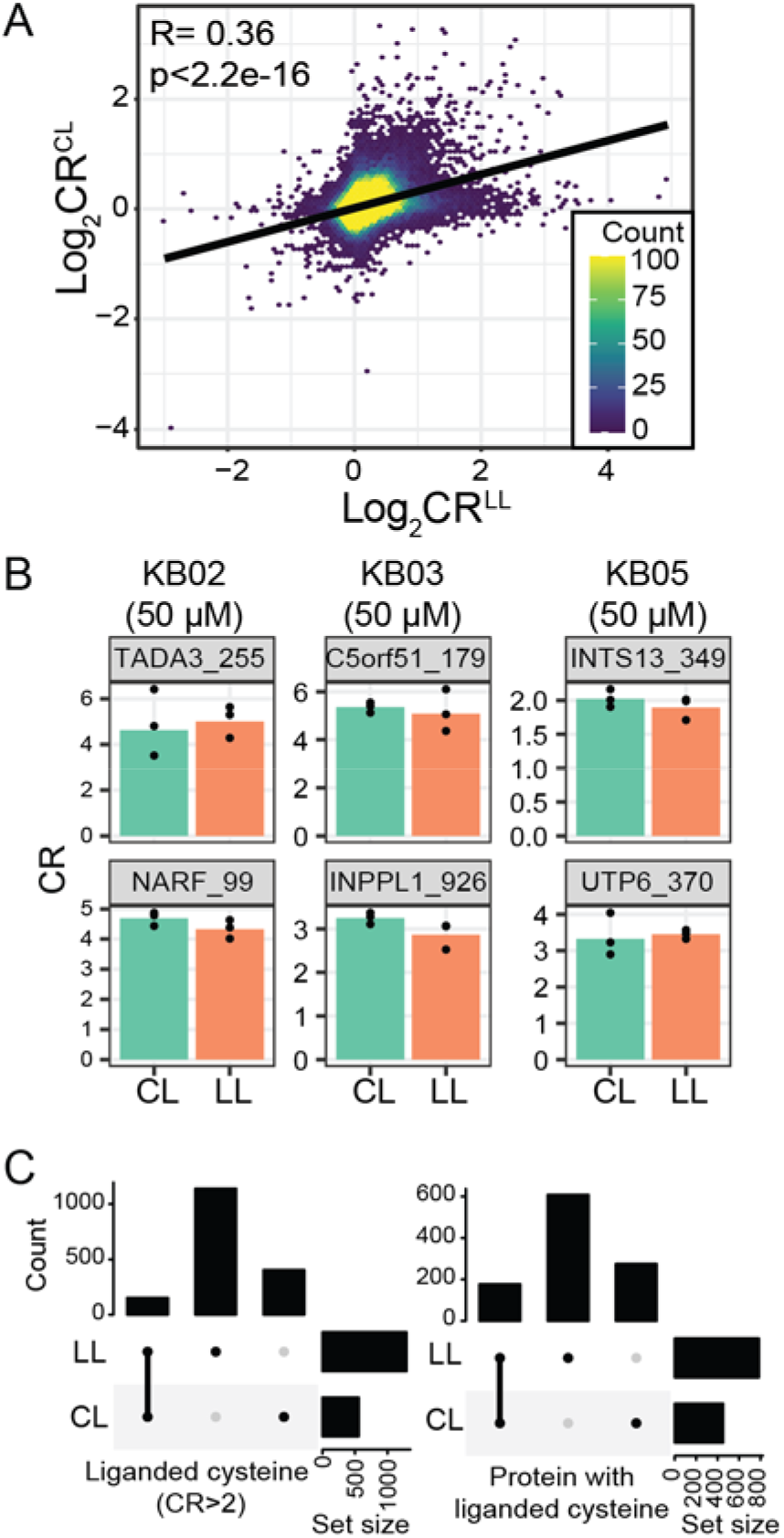
Comparison of cysteine ligandability in CL and LL systems. **A)** Correlation of CRs in two methods. **B)** Examples with consistent CR values in both CL and LL systems. **C)** Upset plot of liganded cysteines (CR > 2) and proteins by at least one probe in each method.

Given that mechanical lysis disrupts cellular membranes, organelles, and transient macromolecular interactions, we postulated that many cysteine residues buried within these higher-order structures in intact cells might become artificially accessible to ligands upon lysis. Consequently, engagement profiles generated in lysates may overrepresent residues that are functionally sequestered in the native cellular environment. We evaluated pPSE scores^33^ for all liganded cysteines (CR>2) in CL and LL datasets. Consistent with this hypothesis, the number of highly exposed cysteines (pPSE < 2) liganded in CL and LL was similar (**Fig. 3A and Fig. S3A**), but LL showed a pronounced increase in liganded cysteines as pPSE increased. In other words, cysteines that are poorly accessible in the native cellular environment are more likely to exhibit ligandability only under lysate conditions. We further extended the comparison to all cysteines under 50 µM KB02 treatment by binning cysteines into three accessibility categories according to their pPSE scores: most accessible (pPSE < 2), moderately accessible (2 < pPSE < 8), and least accessible (pPSE>8). As expected, while obtaining a Pearson R = 0.57 for the most accessible group, the correlation dropped to 0.3 for the least accessible group (**Fig. 3B and Fig. S3B**). We next selected the top differentially liganded (DL) cysteines (Log_2_FC(CR^LL^/CR^CL^) ≥□1 or ≤□-1, and adjust P value < 0.1; **Fig. S2B-C**), and observed consistent patterns across multiple compounds (**Fig. 3C**), with the exception of THZ1. AC19 is excluded from the analysis due to minimal cysteine engagement across both CL and LL conditions and the correspondingly minimal discrepancy between CR^LL^ and CR^CL^ values (**Fig. S2B-C**). This consistency supports that the cysteine accessibility is one major driving factor behind the discrepancy between CL and LL, to a great extent independent of compound identity. Therefore, most DL cysteines correspond to residues with higher pPSE scores and are predicted to be structurally buried (**Fig. 3D**). GO term analysis of these differentially liganded cysteines further revealed similar protein categories across KB02, KB03, KB05, and CL4, despite their different reactivity profiles (**Fig. 3E**). Many of the enriched protein categories correspond to components of the nuclear envelope (**Fig. 3E-F and Fig. S3C**), a structure that is disrupted during lysis. These nuclear envelope cysteines typically exhibit pPSE > 2 and exist in helical structures (**Fig. 3G**). In our experiments, these cysteines were better bound by KB02 (**Fig. 3F**) in lysate than in live cells, with the exception of PCYT1A C73, which is more solvent-exposed and showed higher ligandability in live cells (**Fig. 3G**). The observation is best exemplified by multiple cysteines in XPO1, surface-exposed C528 as the most accessible cysteine, generated higher CR^CL^ than CR^LL^, whereas buried-within-structure cysteines C327 and C585 being less accessible demonstrated reduced ligandability in live cells in comparison with lysate (**Fig. 3H**). Both syringe lysis and sonication are widely used and often interchangeable mechanical lysis methods. To assess whether the choice of lysis method introduces systematic bias in cysteine ligandability measurements, we treated K562 cells with KB02/KB05 and subsequently lysed them by each approach (**Fig. S3D**). In addition to confirming consistent protein abundance profiles (no differential proteins observed; **Fig. S3E**), we observed high correlation in competition ratios (CR) for both KB02 (Pearson r = 0.88) and KB05 (Pearson r = 0.72) (**Fig. S3F**), suggesting that these lysis methods do not significantly bias cysteine engagement results. This consistency was exemplified by XPO1, where all 6 measured cysteines exhibit consistent trends between the two lysis methods (**Fig. S3G**).

**Figure 3.**
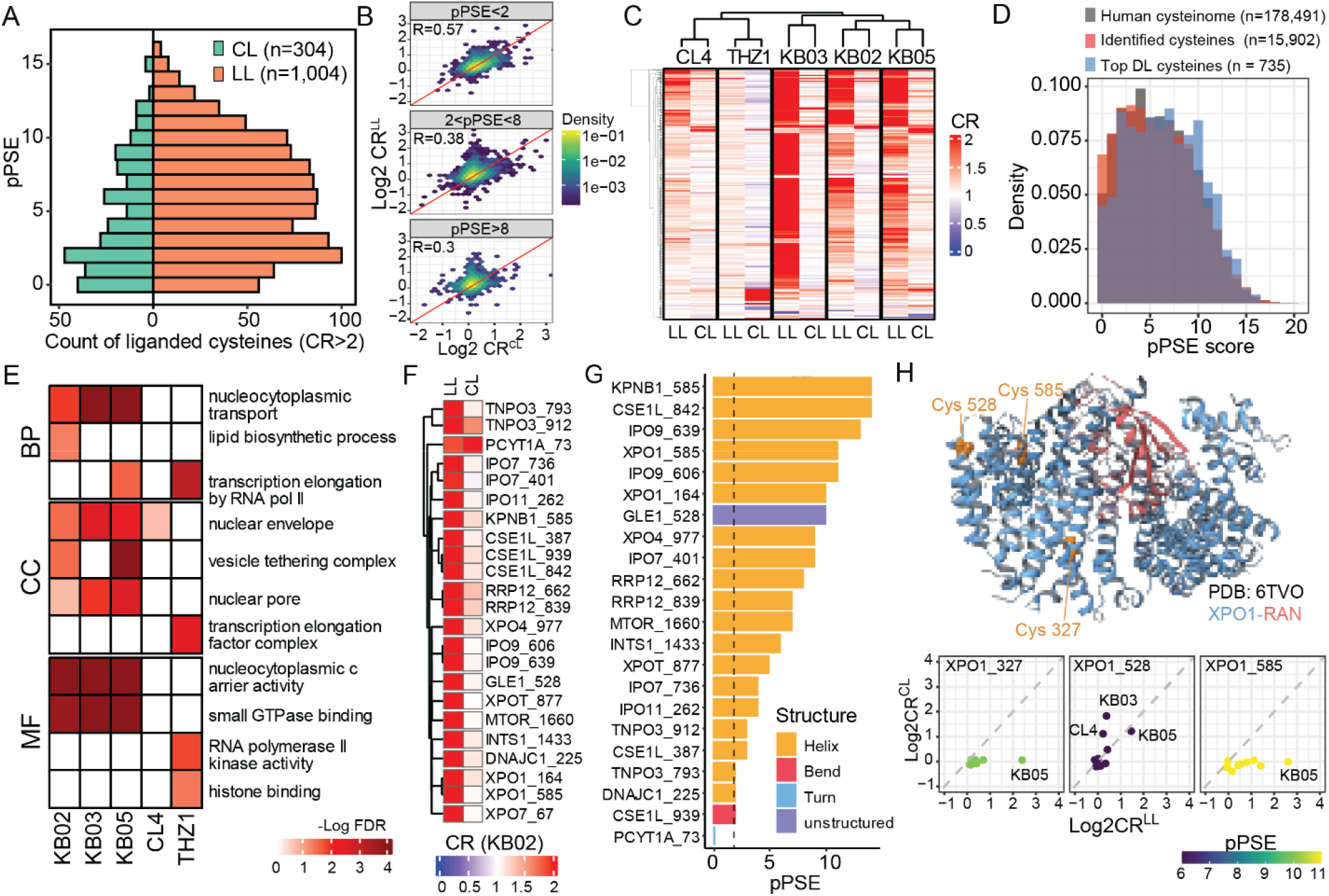
Intrinsic cysteine side chain accessibility as a primary contributor to the discrepancy of cysteine ligandability in cell (CL) and lysate (LL). **A)** Comparison of surface exposure pPSE score of liganded cysteines (CR > 2) in CL and LL. **B)** Comparison of CRs with KB02 in surface-exposed (pPSE ≤2), medium-buried (2 < pPSE ≤ 8) and deep buried (pPSE > 8) cysteine residues. **C)** Heatmap of top differentially liganded (DL) cysteines. **D)** pPSE distribution of all human cysteines, quantified cysteines in this study, and cysteines with different ligandability profiles between CL and LL. **E)** Heatmap of gene ontology (GO) analysis of proteins containing top differentially liganded cysteines. **F)** CRs of top differentially liganded cysteines of nuclear envelope proteins. **G)** pPSE and structural types of cysteines showed in F. Dotted line is pPSE = 2. **H)** Illustration of XPO1 cysteines in protein structure (PDB:6TVO) and their engagement profile in two methods.

Beyond intrinsic cysteine side chain accessibility, our previous work demonstrated that protein abundance change in response to treatment with electrophilic fragments, such as protein degradation, can constitute another confounding factor in interpreting live-cell ABPP data^26^. To more systematically evaluate the effect of protein abundance change, we conducted global proteomics profiling of all drug-treated live cells using flow-through samples from DBIA enrichment. Interestingly, as reflected in the PCA plot, the major separation PC1 was between THZ-1 and all others, with the second PC separating warhead types (acrylamide vs chloroacetamide; **Fig. 4A and Fig. S4A**). THZ1, as a highly selective CDK7 inhibitor, induced the greatest number (∼150) of protein abundance changes (FC > 2, and adjusted P value < 0.05), much higher than the more promiscuous “scout” fragments and less reactive AC19 (**Fig. 4B**)^9^. Before analyzing THZ1-specific protein changes, we first asked to what extent protein abundance changes distort CR^CL^ measurements under the more typical KB02 treatment. Though a previous study by Julio et al. suggests that using excessively high concentration of KB02 (100 µM) may cause proteome remodeling, including aggregation and depletion of the nuclear pore complex^34^, we only observed subtle protein abundance changes using 50 µM KB02 with 20 proteins differentially regulated (FC>2 and adjusted p-value<0.05; **Fig. S4C**). None of the nuclear pore components showed significant abundance changes, further supporting the absence of significant proteome remodeling. We compared protein abundance fold change (DMSO/KB02) with the corresponding CR^CL^ values and observed overlaps, despite only a small fraction (∼1%), between differentially regulated proteins and differentially engaged cysteines (**Fig. 4C and Fig. S4B**). Orange dots in **Fig. 4C** represented cysteines that changed CR^CL^ values without corresponding changes in protein abundance, representing true ligand-cysteine engagement, whereas green dots represented cysteines with concordant changes in CR^CL^ and protein abundance. In addition to ACAT1 C126 as we previously demonstrated^26^, three more ACAT1 cysteines (C119, C196, and C413) showed the similar trend, further corroborating the notion that the observed CR^CL^ values were primarily driven by reduction in protein abundance, although we cannot rule out the possibility that cysteine engagement caused simultaneous protein degradation^34,35^. After correcting CR^CL^ values for protein abundance, three sites of ACAT1 including C119, C126 and C196 reverted to non-significant values (corrected CR^CL^ ≈1) while only C413 remained above the CR threshold, confirming that abundance changes were inflating apparent ligandability (**Fig. 4D**). As another example, the cysteine CR^CL^ of C15orf39 can also be explained by protein abundance change. Our results suggest that apparent CR measurements can be convoluted by protein abundance chance in live cells.

**Figure 4.**
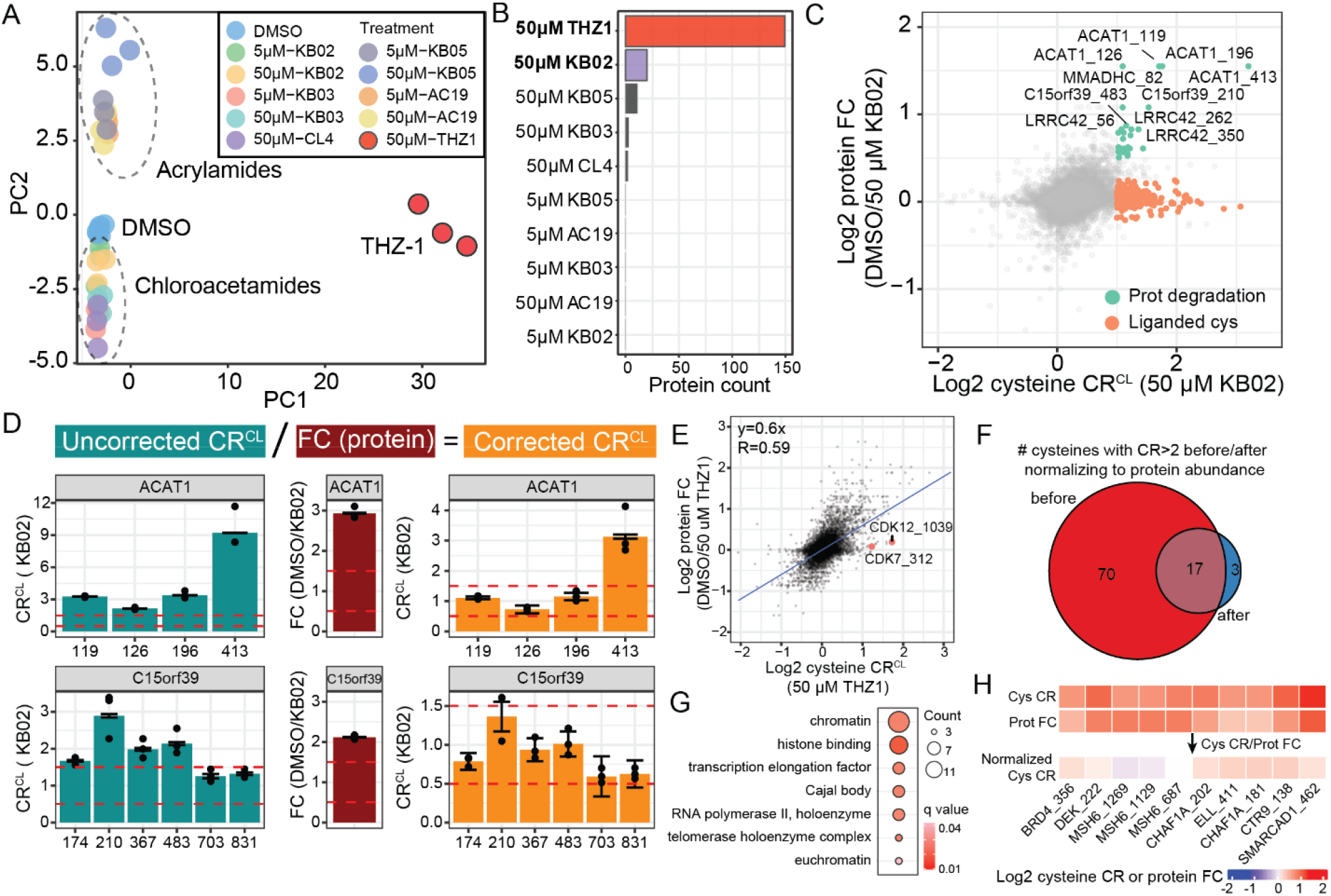
Protein abundance changes in live cell ABPP as a confounding factor. **A)** Principal component analysis (PCA) of global proteomics. 6589 quantified proteins are included. **B)** Number of significantly changed proteins (FC > 2, and adjusted P value < 0.05) by each probe. **C)** Log2-scaled protein FC and cysteine CR^CL^ with KB02. Orange dots are cysteines with Log2 (CR^CL^) > 1 and protein −0.25 < Log2 (protein FC) < 0.25. Green dots are cysteines with Log2 (CR^CL^) > 1 and protein 0.5 < Log2 (protein FC). **D)** Example cysteines from ACAT1 and C15orf39, whose CR^CL^ values reflect reductions in protein abundance induced by KB02. CR^CL^ values are corrected using protein abundance. **E)** Log2-scaled protein FC and cysteine CR^CL^ with THZ1 reveal substantial correlation between protein abundance and cysteine CR^CL^. CDK12_1039 and CDK7_312 are known THZ1 targets and their protein abundances remain constant. **F)** Venn diagram of cysteines with raw CR^CL^ > 2 and protein abundance-corrected CR^CL^ > 2. 53 cysteines (∼76%) are no longer considered liganded after the correction. **G)** GO analysis of the proteins corresponding to the 53 cysteines no longer considered liganded (CR^CL^ > 2) after correcting for protein abundance. **H)** example cysteine CR^CL^, their corresponding protein FCs, and corrected CR^CL^.

When inspecting protein FC and cysteine CR^CL^ with THZ1, an even stronger positive correlation was observed (Pearson R = 0.59; **Fig. 4E**). THZ1 is a selective CDK7 inhibitor, covalently modifying C312, that additionally binds to C1039 on CDK12 and inhibits its activity^31,36^. Consistent with these known engagement events, CDK7 C312 and CDK12 C1039 fell on the x axis, confirming the ligation event. In contrast, many other cysteines fell along the y = x diagonal, indicating that their apparent ligandability was primarily driven by THZ1-induced changes in protein abundance rather than direct cysteine modification (**Fig. 4E**). Of the 70 cysteines with raw CR^CL^ > 2, 53 were normalized away after accounting for protein abundance (**Fig. 4F**). Intriguingly, these cysteines were highly enriched on proteins that have histone or chromatin binding activity (**Fig. 4G, H**), leading us to hypothesize that THZ1 promoted many proteins to bind to chromatin and therefore were lost during sample preparation when insoluble proteins and cell debris were removed by mild centrifugation (1,400 RCF for 5 mins)^37^.

To test whether more proteins became associated with chromatin upon THZ1 treatment, we performed subcellular proteomics analysis of chromatin-associated and cytoplasmic proteins after 30- and 60-minute THZ1 treatment (**Fig. 5A, B**)^38,39^. THZ1 inhibits CDK7, which phosphorylates the C-terminal repeat domain (CTD) of RNA polymerase II, and thereby halting transcription^31^. As a result, components of the pre-initiation complex, including GTF2F1 and GTF2F2, became trapped on chromatin (**Fig. 5C and S5**). CDK7 activity is required to recruit negative elongation factors to the promoter-proximal pausing site. As expected, its inhibition released these factors (NELFA, NELFB, and NELFE in **Fig. 5C**)^38,40^. Additionally, we observed a reduction of several CTD phosphorylation-dependent splicing factors, including SCAF1, SCAF4, SCAF11 and SCAF8 (**Fig. 5C**), consistent with previous findings^38^. Another early transcription elongation factor, RTF1, that regulates transcriptional pausing^41^, also accumulated at the pausing site (**Fig. 5D**).

**Figure 5.**
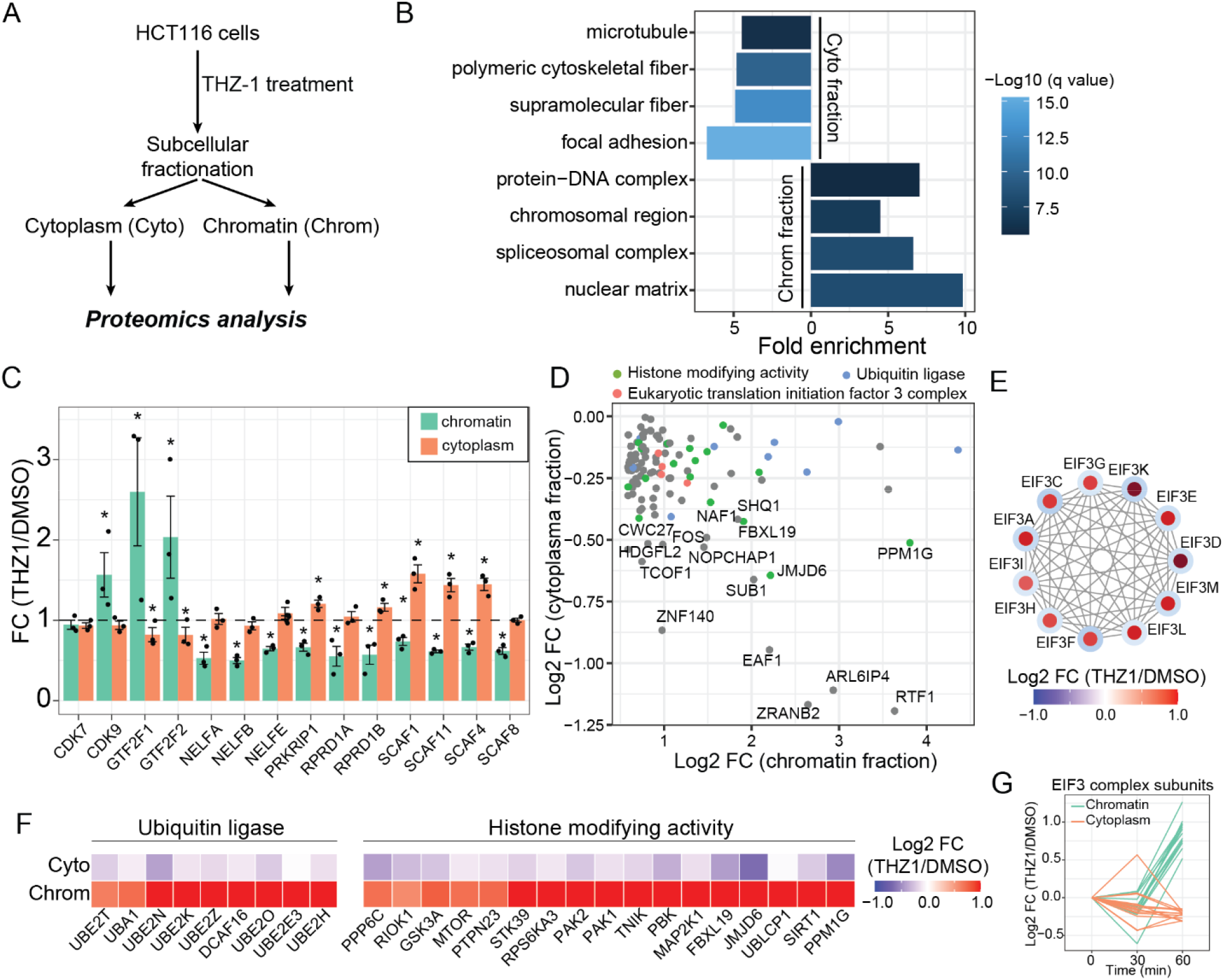
THZ-1 induces protein relocalization. **A)** Workflow for subcellular proteomics used to measure protein relocalization upon THZ1 treatment. **B)** Representative GO terms enriched from top 100 proteins in each subcellular fraction, ranked by number of peptide spectrum matches (PSMs). **C)** Known proteins that relocalize to or away from chromatin upon THZ-1 treatment show the expected changes. * indicates Student’s t-test p < 0.05. **D)** Proteins enriched in the chromatin fraction after THZ1 treatment. Proteins with histone modifying activity, involved in ubiquitin ligase system, and belonging to the translation initiation factor 3 complex are highlighted. **E)** Interaction network of the quantified EIF3 complex subunits. Edges represent physical interactions. Node colors represent log2 FC (THZ1/DMSO) in the chromatin fraction. Border colors represent log2 FC (THZ1/DMSO) in the cytoplasmic fraction. **F)** Fold changes of proteins with ubiquitin ligase activity and histone modifying activity as colored in D. **G)** Fold changes of EIF3 complex subunits at 30 and 60 minutes post-THZ1 treatment.

Beyond proteins with well-established roles in the regulation of early transcriptional pausing, we found rapid and increased chromatin retention of many other proteins, including ubiquitin ligases (e.g. UBE2O, DCAF16), histone-modifying enzymes (e.g. PAK1/2), and several subunits of the eukaryotic initiation factor 3 complex (**Fig. 5D-F**). The accumulation of ubiquitin ligases may reflect a mechanism to alleviate disrupted transcription machinery, as a previous study linked CDK7 inhibition by THZ1 with enhanced degradation of the Pol II complex, though the responsible ligases remain elusive^42^. The ligases identified here therefore are potential candidates pending further investigation. Likewise, many histone modifying enzymes, including kinase MTOR through its noncanonical function^43^, demethylase JMJD6^44^, and deacetylase SIRT1^45^, relocated to chromatin in response to altered transcription pause-release dynamics. We were particularly intrigued by the rapid accumulation of EIF3 complex after only 30 minutes. No accumulation was detected prior to 30 minutes (**Fig. 5G**), consistent with the report from Velychok and colleagues^38^. Beyond their primary cytosolic location and function in translation, various reports suggest the nuclear localization and chromatin association of EIF3 subunits to execute less well-studied functions such as DNA damage response, protein degradation, and mRNA processing^46–50^. We posit that the THZ1-induced accumulation of EIF3 on chromatin reflects its association with nascent RNA molecules that become trapped at stalled Pol II complexes during acute transcriptional shutdown.

Our observation suggests that the mild centrifugation to extract native and soluble proteome for in vitro assays, a common practice in research^51–54^, can lead to misleading ABPP readouts due to the loss of chromatin- and DNA-binding proteins. We decided to test whether omitting the removal of insoluble proteomes in our live cell ABPP workflow would mitigate these artifacts. We performed global proteomic analysis in addition to ABPP using the whole proteome containing both insoluble and soluble fractions (**Fig. 6A**). Those cysteines that previously appeared to have significant competition ratios due to enhanced chromatin binding of their parent proteins (**Fig. 4H**) were no longer significant (grey dots in **Fig. 6B**), consistent with their stable protein abundance when both soluble and insoluble fractions were retained (**Fig. 6C**). As a result, only minimal correlation was observed between protein FC and cysteine CR, indicative of minimal convolution from protein abundance (**Fig. 6D**). As THZ1 is a potent sub-micromolar inhibitor^22,31^, we further investigated whether chromatin relocalization observed at 50 µM could be recapitulated at lower doses (**Fig. S6A**). We confirmed that CDK7 C312 and CDK12 C1039 were engaged by THZ1 in a dose-dependent manner (**Fig. S6B**). 50% occupancy of CDK7 C312 (CR>2) was achieved at 240 nM THZ1 whereas off-target cysteines, such as ITPKA C342 and ARL6IP4 C36, required concentrations of 7 µM or higher to reach equivalent occupancy (**Fig. S6C**). Furthermore, analysis of the significantly altered sub-proteome (adjusted ANOVA p < 0.001) revealed that numerous proteins underwent relocalization between the cytosol and chromatin despite negligible changes in total protein abundance (**Fig. S6D**).

**Figure 6.**
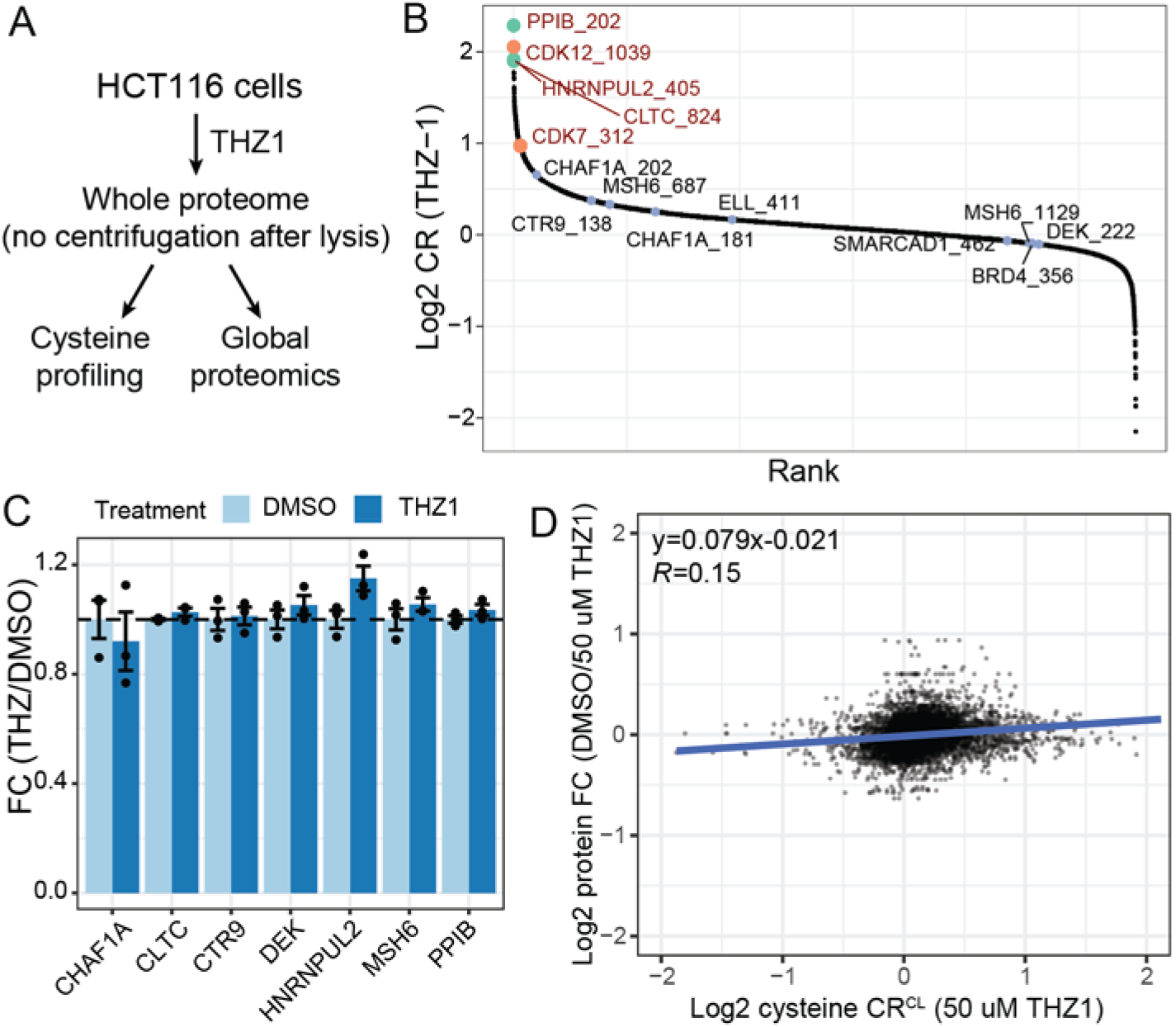
Live-cell ABPP using whole proteome without centrifugation after lysis. **A)** Workflow to analyze cysteine and protein abundance using live HCT116 cells treated with THZ-1. No centrifugation is performed after lysis to retain both soluble and insoluble proteome fractions. **B)** Ranked cysteine CR^CL^ values. Top liganded and known targets are colored green and orange, respectively. Cysteine residues in Figure 4H from chromatin binding proteins are colored grey and no longer show significant engagement. **C)** No alteration in protein abundance of chromatin binding proteins in Figure 4H. **D)** Minimal correlation between protein FC and cysteine CR^CL^ is observed compared to Figure 4E.

In summary, our systematic comparison of live cell- and lysate-based TMT-ABPP reveals that discrepancies in measured cysteine ligandability arise from a set of biologically interpretable factors. Predicted cysteine accessibility is the dominant determinant of differential engagement between systems, particularly for residues buried within membranes, organelles, or higher-order protein assemblies that are disrupted upon lysis. In live cells, these effects are further compounded by changes in protein abundance, such as KB02-induced protein degradation, and protein relocalization to insoluble fractions, as observed upon THZ1 treatment where proteins associate with chromatin and are consequently depleted during sample handling. Importantly, we show that standard sample preparation practices, such as removal of insoluble cell debris, can lead to misinterpretation of apparent cysteine engagement. Together, these findings establish a framework for selecting ABPP platforms and interpreting results based on underlying biology, and underscore the importance of accounting for intrinsic residue accessibility, protein abundance, and protein localization when designing live-cell cysteine profiling experiments. We recommend performing large-scale profiling of cysteine-targeting compounds in the native full proteome lysate to maximize throughput (e.g., 96-well format) and reproducibility while minimizing potential biases introduced by complex cellular processes, unless full cellular contexts and secondary responses are desired and accounted for with orthogonal experiments. Once hit or lead compounds are identified, their target engagement should be validated in a cellular context. For these cellular ABPP assays, we suggest using shorter incubation times and lower concentrations to mitigate proteome-wide remodeling. It is important to note that the common centrifugation step used to clear lysate should be omitted when possible, as proteins relocalized into insoluble fractions (e.g., chromatin) are lost during sedimentation, generating false positive cysteine engagement signals that reflect protein loss rather than direct ligand-cysteine interaction. In addition, changes in protein abundance must be accounted for in follow-up analyses by either global proteomics or other types of protein assays such as immunoblot or ELISA to support mechanistic interpretation. We also provide practical solutions to address confounding effects due to protein abundance change and protein relocalization. More broadly, we anticipate that these considerations apply not only to reactive-cysteine profiling but to all ABPP approaches that measure amino acid modification states (e.g., tyrosine, serine). Given that competitive ABPP relies on indirect loss-of-signal readouts, many of the discrepancies we observe, including those arising from abundance changes, or relocalization, highlight the value of an orthogonal gain-of-signal validation strategy. A targeted measurement of compound-cysteine adduct ions would provide direct evidence of covalent modification and help distinguish true ligandable sites from apparent CR changes driven by indirect biological effects. In addition, inspired by our observation, we anticipate the subcellular fraction can be further expanded to subcellular ABPP to specifically identify molecules that induce protein trafficking across compartments.

## Methods

### Chemicals

KB02 (cat. # 911798), KB03 (cat. # 912654) and KB05 (cat. # 912131) were purchased from Sigma-Aldrich. THZ-1 (cat. # S7549), was purchased from Selleckchem. CL4 and AC19 was obtained from a customized library and DBIA probe was synthesized as described in previous publication^23^.

### Cell culture

K562 cells were grown in RMPI-1640 (Corning) medium supplemented with 10% FBS and 1 % Penicillin/Streptomycin. HCT116 cells were grown in DMEM (Corning) medium supplemented with 10% FBS and 1 %. All cell lines were cultured in 5% CO_2_ at 37 °C.

### In-lysate cysteine profiling

For in-lysate cysteine profiling, cells at ∼80% confluency were collected, washed twice with cold PBS, and stored at −80□°C until lysis. Cysteine profiling was performed as previously reported^19,23^. Frozen cell pallet was resuspended in lysis buffer (PBS, pH 7.4, 0.1% NP-40) and lysed by passing through a 21-gauge needle followed by sonication (3s-on, 3s-off, 50% amplitude). Soluble fraction was collected after centrifugation at 1,400 g for 5 minutes while whole proteome was collected without centrifugation. Cell lysate (soluble fraction or whole proteome) was diluted with lysis buffer and 15 µL lysate (2 µg/µL protein) was dispensed into each well of a 96-well plate. Compound solution prepared in lysis buffer was added to each well to a final concentration of 50□µM and incubated for 1□h at room temperature.

### In-cell cysteine profiling

For in-cell cysteine profiling, 5 × 10^5^ cells were seeded in 12-well plate and then treated with compound for 1 hour at 37 °C before cells were collected as described above. Frozen cell pellets were resuspended in lysis buffer (PBS, pH 7.4, 0.1% NP-40) and lysed by passing through a 21-gauge needle. Clear cell lysate was collected after centrifugation at 1,400 g for 5 minutes while whole proteome lysate (related to Fig. 6) was not centrifugated. Protein concentration was measured by BCA assay.

### DBIA labeling and SP3 cleanup

For both live cell and lysate workflows, DBIA was added in lysis buffer to a final concentration of 500□µM and incubated in the dark for 1□h. 3 µL SP3 beads (1:1 mixture of hydrophobic and hydrophilic type) was added to 20 µg proteins followed by precipitation buffer (20 mM DTT in ∼98% ethanol) to reach final >50% ethanol. Samples were incubated for 15□min with gentle shaking, placed on a magnetic stand, and the supernatant was aspirated. Beads were washed once with 80% ethanol, resuspended in 30□µL alkylation buffer (20□mM IAA in lysis buffer), and incubated in the dark for 30□min. An additional 60□µL precipitation buffer was added, and SP3 washes were performed as previously described (three washes)^23,55^. Beads were then resuspended in 20□µL of 200□mM EPPS buffer (pH□8.5) containing 0.3□µg Lys-C and incubated for 3□h at room temperature. Afterward, 20□µL EPPS buffer containing 0.3□µg trypsin was added, and digestion proceeded overnight at 37□°C.

### TMT labeling and enrichment

The next day, acetonitrile and TMTpro reagent (50 µg; TMT:peptide = 2.5:1, w/w) were sequentially added to the peptide-bead mixture to achieve a final acetonitrile concentration of ∼30% (v/v). The reaction was gently mixed for 60 min at room temperature. Reaction was quenched by adding 5% hydroxyl amine to final concentration at 0.5%. TMT-labeled peptides from each sample were combined, dried by SpeedVac, and desalted using a 100-mg Sep-Pak column.

Desalted TMT-labeled peptides were resuspended in 460 µL100 mM HEPES buffer (pH 7.4). Streptavidin enrichment was performed by adding 80□µL Pierce-High Capacity Streptavidin Agarose (Cat.□#20359) and rotating for 3□h at room temperature. The peptide-bead mixture was transferred to an Ultrafree-MC centrifugal filter (hydrophilic PTFE, 0.22□µm) and centrifuged at 1,000□g for 30□s. The flow-through was retained for protein abundance analysis. Beads were washed twice with wash buffer 1(100 mM HEPES, pH 7.4, 0.05% NP-40), three times with wash buffer 2 (100 mM HEPES, pH 7.4) and once with water. Peptides were eluted twice with elution buffer (80% acetonitrile, 0.1% formic acid) at room temperature for 10□min each, followed by two additional elutions at 72□°C for 10□min each. Combined eluates were dried in a SpeedVac and desalted using StageTips prior to LC-FAIMS-hrMS^2^ analysis.

### Subcellular fractionation

Subcellular fractionation of cytosolic and chromatin proteome was performed as previously reported^38,39^. Briefly, cell pellet was collected after compound treatment, resuspended in lysis buffer 1 (PBS, pH 7.4, 0.1% NP-40), and lysed by gentle pipetting, followed by incubation for 5 minutes on ice. The lysate was carefully overlaid onto buffer□1 containing 24% sucrose in PBS by slowly pipetting along the wall of a LoBind tube. After centrifugation at 3,500□×□g for 10□min, the pellet (nuclear fraction) was collected, and the supernatant was transferred to a new tube. The cytosolic fraction was collected from the clarified supernatant after centrifugation at 14,000□×□g for 1□min. The nuclei fraction was further fractionated by resuspending the nuclei pellet in buffer 2 (50% glycerol in PBS) and then immediately adding lysis buffer 2 (PBS, 1M urea, 1% NP40). The mixture was vortexed briefly (4□s) and incubated on ice for 2□min, then centrifuged at 14,000□×□g for 2□min. The precipitated pellet was briefly rinsed with PBS, resuspended in lysis buffer 3 (200 mM EPPS, pH = 8.5, 8M urea) and collected as chromatin fraction. All fractions were immediately frozen and stored at −80□°C until processing.

### Global proteomics analysis and HPLC Fractionation

Full or subcellular proteome containing ∼30 µg proteins was directly loaded to SP3 beads for clean-up, reduction/alkylation and TMT-labeling as described above. ∼ 100 µg TMT-labeled flow-through peptides from cysteine profiling or whole or subcellular proteomes were resuspended in water and fractionated using basic pH reversed-phase (BPRP) HPLC^56^. The resulting 96 fractions were consolidated into 12 or 24 fractions, which were then desalted and analyzed by FAIMS-hrMS^2^ analysis.

### LC-FAIMS-hrMS^2^ analysis

Samples were resuspended in LC-MS loading buffer (5% ACN and 5% FA) and loaded on a 100-µm capillary column packed with 30 cm of Accucore 150 resin (2.6 □m, 150Å; Thermo Fisher Scientific). Enriched cysteine and full proteome samples were separated using a 180-min and 90-min methods, respectively, on a Proxeon NanoLC-1200 UPLC system. Data were collected using a high-resolution MS2 method on an Orbitrap Eclipse mass spectrometer coupled with a FAIMS Pro device. Data were collected alternating between a set of three FAIMS compensation voltages (CVs). For enriched cysteine sample, two sets of CV values (−60, −45 and −35V; −70, −55 and −30) were used for double-shot analysis^23^. For full proteome analysis, only one set of CV values (−80, −60 and −40V) was used for single-shot analysis. MS1 scans were collected in the Orbitrap with a resolution setting of 60 K, a mass range of 400-1600 m/z, an AGC at 100%, and a maximum injection time of 50 ms. MS2 scans were acquired in Top Speed mode with a cycle time of 1 s. Peptide precursors were selected and fragmented using HCD with a collision energy of 36. MS2 scans were collected in the Orbitrap with a resolution of 50K, a fixed scan range of 110-2000 m/z, and a 500% AGC with a maximum injection time of 86 ms. Dynamic exclusion was set to 120 s for cysteine and 90 s for full proteome with a mass tolerance of ± 10 p.p.m.. Unfractionated flow-through from each cysteine sample was separated using a 60-min method and analyzed as full proteome analysis for normalization of cysteine profiling.

### Proteomic Data analysis

Raw files were searched using the Comet search engine (ver. 2019.01.5)^57^ with the UniProt human proteome database (downloaded 11/24/2021) with contaminants and reverse decoy sequences appended. Precursor error tolerance was 50 p.p.m. and fragment error tolerance was 0.02 Da. Static modifications included Cys carboxyamidomethylation (+57.0215) and TMTpro (+304.2071) on Lys and peptide N-termini. Methionine oxidation (+15.9949) was allowed as variable modification. DBIA-modification on cysteine residues (+239.1628) was set as variable modification for searching cysteine profiling files. Peptide spectral matches were filtered to a peptide false discovery rate (FDR) of <1% using linear discriminant analysis employing a target-decoy strategy^58,59^. Resulting peptides were further filtered to obtain a 1% protein FDR at the entire dataset level (including all plexes in an experiment)^60^. Cysteine-modified peptides were site-localized using AScorePro, with a cutoff of 13 (P < 0.05)^61,62^. For quantification of each MS2 spectrum, reporter ion signal-to-noise (S/N) were adjusted to correct for impurities during synthesis of different TMT reagents according to the manufacturer’s specifications. A sum S/N values of all reporter ions of >100 was required. Protein quantification was normalized by the sum of the S/N for all proteins in each channel to correct for protein loading. Similarly, cysteine site quantification was normalized. Proteins or cysteine sites were quantified by summing the reporter-ion S/N values across all peptide-spectrum matches. pPSE scores of cysteines with confident AlphaFold2 prediction (pLDDT>70) were obtained from literature^33^ and applied to annotate the cysteines identified in this work to evaluate their side-chain accessibility.

## Statistical analysis

All analysis was conducted by R software (version 4.5.1). All P values were calculated by either t-test or one-way ANOVA and then adjusted by Benjamini-Hochberg method to correct for multiple hypothesis testing. Gene ontology and other pathway analysis was performed by ClusterProfiler 4.0^63^. Protein-protein interaction and network analysis was performed and graphed using Cytoscape^64^.

## Acknowledgments

This work was funded in part by NIH grants GM67945 (S.P.G.), CA282268 (Q.Y.), Worcester Foundation for Biomedical Research (Q.Y.), and ASMS Research Award (Q.Y.).

## Supporting Information

- Competition ratio distribution across two systems under different drug treatments (Figure S1); Comparison of cysteine ligandability across two systems under different drug treatments (Figure S2); Comparison of cysteine pPSE distribution (Figure S3); Protein abundance changes induced by compound treatment in live cells (Figure S4); Subcellular proteomics with THZ1 (Figure S5); Subcellular proteomics across a THZ1 dose range (Figure S6).
- TMT-ABPP cysteine profiling in native proteome with chemical probes in-cell vs. in-lysate treatment (Table S1); TMT profiling of full proteome following in-cell chemical probes treatment (Table S2); TMT profiling of subcellular proteomes from THZ-1 in-cell treatment (Table S3); TMT-ABPP cysteine profiling in full and chromatin proteomes from THZ1 in-cell treatment (Table S4); TMT-ABPP cysteine profiling comparing syringe lysis and sonication using in-cell scout probe treatment (Table S5); TMT proteomics comparing syringe lysis and sonication using in-cell scout probe treatment (Table S6); TMT-ABPP cysteine profiling in full proteome, THZ1 dose-response, in-cell treatment (Table S7); TMT profiling of subcellular proteomes, THZ1 dose-response, in-cell treatment (Table S8).

## References

(1) Cravatt, B. F.; Wright, A. T.; Kozarich, J. W. Activity-Based Protein Profiling: From Enzyme Chemistry to Proteomic Chemistry. Annu. Rev. Biochem. 2008, 77 (Volume 77, 2008), 383–414.

(2) Niphakis, M. J.; Cravatt, B. F. Ligand Discovery by Activity-Based Protein Profiling. Cell Chem. Biol. 2024, 31 (9), 1636–1651.

(3) Nomura, D. K.; Dix, M. M.; Cravatt, B. F. Activity-Based Protein Profiling for Biochemical Pathway Discovery in Cancer. Nat. Rev. Cancer 2010, 10 (9), 630–638.

(4) Fonovic, M.; Bogyo, M. Activity Based Probes for Proteases: Applications to Biomarker Discovery,Molecular Imaging and Drug Screening. http://www.eurekaselect.com.

(5) Waxman, D. J.; Strominger, J. L. Cephalosporin-Sensitive Penicillin-Binding Proteins of Staphylococcus Aureus and Bacillus Subtilis Active in the Conversion of [14C]Penicillin G to [14C]Phenylacetylglycine. J. Biol. Chem. 1979, 254 (23), 12056–12061.

(6) Jessani, N.; Niessen, S.; Wei, B. Q.; et al. A Streamlined Platform for High-Content Functional Proteomics of Primary Human Specimens. Nat. Methods 2005, 2 (9), 691–697.

(7) Weerapana, E.; Wang, C.; Simon, G. M.; et al. Quantitative Reactivity Profiling Predicts Functional Cysteines in Proteomes. Nature 2010, 468 (7325), 790–795.

(8) Fonovi□, M.; Bogyo, M. Activity Based Probes for Proteases: Applications to Biomarker Discovery, Molecular Imaging and Drug Screening. Curr. Pharm. Des. 2007, 13 (3), 253–261.

(9) Backus, K. M.; Correia, B. E.; Lum, K. M.; et al. Proteome-Wide Covalent Ligand Discovery in Native Biological Systems. Nature 2016, 534 (7608), 570–574.

(10) Zhang, X.; Crowley, V. M.; Wucherpfennig, T. G.; et al. Electrophilic PROTACs That Degrade Nuclear Proteins by Engaging DCAF16. Nat. Chem. Biol. 2019, 15 (7), 737–746.

(11) Brulet, J. W.; Ciancone, A. M.; Yuan, K.; et al. Advances in Activity-Based Protein Profiling of Functional Tyrosines in Proteomes. Isr. J. Chem. 2023, 63 (3–4), e202300001.

(12) Abbasov, M. E.; Kavanagh, M. E.; Ichu, T.-A.; et al. A Proteome-Wide Atlas of Lysine-Reactive Chemistry. Nat. Chem. 2021, 13 (11), 1081–1092.

(13) Nemmara, V. V.; Thompson, P. R. Development of Activity-Based Proteomic Probes for Protein Citrullination. In Activity-Based Protein Profiling; Cravatt, B. F., Hsu, K.-L., Weerapana, E., Eds.; Springer International Publishing: Cham, 2019; pp 233–251.

(14) Xiao, H.; Jedrychowski, M. P.; Schweppe, D. K.; et al. A Quantitative Tissue-Specific Landscape of Protein Redox Regulation during Aging. Cell 2020, 180 (5), 968–983.

(15) Takahashi, M.; Chong, H. B.; Zhang, S.; et al. DrugMap: A Quantitative Pan-Cancer Analysis of Cysteine Ligandability. Cell 2024, 187 (10), 2536-2556.e30.

(16) Yang, F.; Jia, G.; Guo, J.; et al. Quantitative Chemoproteomic Profiling with Data-Independent Acquisition-Based Mass Spectrometry. J. Am. Chem. Soc. 2022, 144 (2), 901–911.

(17) Boatner, L. M.; Palafox, M. F.; Schweppe, D. K.; et al. CysDB: A Human Cysteine Database Based on Experimental Quantitative Chemoproteomics. Cell Chem. Biol. 2023, 30 (6), 683-698.e3.

(18) Biggs, G. S.; Cawood, E. E.; Vuorinen, A.; et al. Robust Proteome Profiling of Cysteine-Reactive Fragments Using Label-Free Chemoproteomics. Nat. Commun. 2025, 16 (1), 73.

(19) He, Y.; Yang, K.; Li, S.; et al. TMT-Based Multiplexed (Chemo)Proteomics on the Orbitrap Astral Mass Spectrometer. Mol. Cell. Proteomics 2025, 24 (5), 100968.

(20) Wang, Y.; Dix, M. M.; Bianco, G.; et al. Expedited Mapping of the Ligandable Proteome Using Fully Functionalized Enantiomeric Probe Pairs. Nat. Chem. 2019, 11 (12), 1113–1123.

(21) Li, M.; Patel, H. V.; Cognetta, A. B.; et al. Identification of Cell Wall Synthesis Inhibitors Active against Mycobacterium Tuberculosis by Competitive Activity-Based Protein Profiling. Cell Chem. Biol. 2022, 29 (5), 883-896.e5.

(22) Kuljanin, M.; Mitchell, D. C.; Schweppe, D. K.; et al. Reimagining High-Throughput Profiling of Reactive Cysteines for Cell-Based Screening of Large Electrophile Libraries. Nat. Biotechnol. 2021, 39 (5), 630–641.

(23) Yang, K.; Whitehouse, R. L.; Dawson, S. L.; et al. Accelerating Multiplexed Profiling of Protein-Ligand Interactions: High-Throughput Plate-Based Reactive Cysteine Profiling with Minimal Input. Cell Chem. Biol. 2024, 31 (3), 565-576.e4.

(24) Thompson, A.; Schäfer, J.; Kuhn, K.; et al. Tandem Mass Tags:□ A Novel Quantification Strategy for Comparative Analysis of Complex Protein Mixtures by MS/MS. Anal. Chem. 2003, 75 (8), 1895–1904.

(25) Li, J.; Van Vranken, J. G.; Pontano Vaites, L.; et al. TMTpro Reagents: A Set of Isobaric Labeling Mass Tags Enables Simultaneous Proteome-Wide Measurements across 16 Samples. Nat. Methods 2020, 17 (4), 399–404.

(26) Dong, K. D.; Yu, Q.; Yang, K.; et al. Enrichment-Free, Targeted Covalent Drug Discovery in Live Cells. ACS Chem. Biol. 2025, 20 (12), 2851–2862.

(27) Vinogradova, E. V.; Zhang, X.; Remillard, D.; et al. An Activity-Guided Map of Electrophile-Cysteine Interactions in Primary Human T Cells. Cell 2020, 182 (4), 1009-1026.e29.

(28) Njomen, E.; Hayward, R. E.; DeMeester, K. E.; et al. Multi-Tiered Chemical Proteomic Maps of Tryptoline Acrylamide–Protein Interactions in Cancer Cells. Nat. Chem. 2024, 16 (10), 1592–1604.

(29) Zhang, C.; Zhou, C.; Magassa, A.; et al. A Platform for Mapping Reactive Cysteines within the Immunopeptidome. Nat. Commun. 2024, 15 (1), 9698.

(30) Bergen, W. van; Hevler, J.F.; Wu, W.; et al. Site-Specific Activity-Based Protein Profiling Using Phosphonate Handles. Mol. Cell. Proteomics 2023, 22 (1).

(31) Kwiatkowski, N.; Zhang, T.; Rahl, P. B.; et al. Targeting Transcription Regulation in Cancer with a Covalent CDK7 Inhibitor. Nature 2014, 511 (7511), 616–620.

(32) Bludau, I.; Willems, S.; Zeng, W.-F.; et al. The Structural Context of Posttranslational Modifications at a Proteome-Wide Scale. PLOS Biol. 2022, 20 (5), e3001636.

(33) White, M. E. H.; Gil, J.; Tate, E. W. Proteome-Wide Structural Analysis Identifies Warhead- and Coverage-Specific Biases in Cysteinefocused Chemoproteomics. Cell Chem. Biol. 2023, 30 (7), 828-838.e4.

(34) Julio, A. R.; Shikwana, F.; Truong, C.; et al. Delineating Cysteine-Reactive Compound Modulation of Cellular Proteostasis Processes. Nat. Chem. Biol. 2025, 21 (5), 693–705.

(35) King, E. A.; Cho, Y.; Hsu, N. S.; et al. Chemoproteomics-Enabled Discovery of a Covalent Molecular Glue Degrader Targeting NF-□B. Cell Chem. Biol. 2023, 30 (4), 394-402.e9.

(36) Zhang, T.; Kwiatkowski, N.; Olson, C. M.; et al. Covalent Targeting of Remote Cysteine Residues to Develop CDK12 and CDK13 Inhibitors. Nat. Chem. Biol. 2016, 12 (10), 876–884.

(37) dos Santos, B.; Bion, M. C.; Goujon-Svrzic, M.; et al. REAP+: A Single Preparation for Rapid Isolation of Nuclei, Cytoplasm, and Mitochondria. Anal. Biochem. 2024, 687, 115445.

(38) Velychko, T.; Mohammad, E.; Ferrer-Vicens, I.; et al. CDK7 Kinase Activity Promotes RNA Polymerase II Promoter Escape by Facilitating Initiation Factor Release. Mol. Cell 2024, 84 (12), 2287-2303.e10.

(39) Conrad, T.; Ørom, U. A. Cellular Fractionation and Isolation of Chromatin-Associated RNA. In Enhancer RNAs: Methods and Protocols; Ørom, U. A., Ed.; Springer: New York, NY, 2017; pp 1–9.

(40) Larochelle, S.; Amat, R.; Glover-Cutter, K.; et al. Cyclin-Dependent Kinase Control of the Initiation-to-Elongation Switch of RNA Polymerase II. Nat. Struct. Mol. Biol. 2012, 19 (11), 1108–1115.

(41) Langenbacher, A. D.; Lu, F.; Tsang, L.; et al. Rtf1-Dependent Transcriptional Pausing Regulates Cardiogenesis. eLife 2026, 13.

(42) Sun, Y.; Zhang, Y.; Schultz, C. W.; et al. CDK7 Inhibition Synergizes with Topoisomerase I Inhibition in Small Cell Lung Cancer Cells by Inducing Ubiquitin-Mediated Proteolysis of RNA Polymerase II. Mol. Cancer Ther. 2022, 21 (9), 1430– 1438.

(43) Zhao, T.; Fan, J.; Abu-Zaid, A.; et al. Nuclear mTOR Signaling Orchestrates Transcriptional Programs Underlying Cellular Growth and Metabolism. Cells 2024, 13 (9).

(44) Liu, W.; Ma, Q.; Wong, K.; et al. Brd4 and JMJD6-Associated Anti-Pause Enhancers in Regulation of Transcriptional Pause Release. Cell 2013, 155 (7), 1581–1595.

(45) Zhang, T.; Kraus, W. L. SIRT1-Dependent Regulation of Chromatin and Transcription: Linking NAD+ Metabolism and Signaling to the Control of Cellular Functions. Biochim. Biophys. Acta 2010, 1804 (8), 1666–1675.

(46) Kachaev, Z. M.; Ivashchenko, S. D.; Kozlov, E. N.; et al. Localization and Functional Roles of Components of the Translation Apparatus in the Eukaryotic Cell Nucleus. Cells 2021, 10 (11), 3239.

(47) Sha, Z.; Brill, L. M.; Cabrera, R.; et al. The eIF3 Interactome Reveals the Translasome, a Supercomplex Linking Protein Synthesis and Degradation Machineries. Mol. Cell 2009, 36 (1), 141–152.

(48) Watkins, S. J.; Norbury, C. J. Cell Cycle-Related Variation in Subcellular Localization of eIF3e/INT6 in Human Fibroblasts. Cell Prolif. 2004, 37 (2), 149–160.

(49) Yen, H.-C. S.; Chang, E. C. Yin6, a Fission Yeast Int6 Homolog, Complexes with Moe1 and Plays a Role in Chromosome Segregation. Proc. Natl. Acad. Sci. 2000, 97 (26), 14370–14375.

(50) Shi, Y.; Giammartino, D. C. D.; Taylor, D.; et al. Molecular Architecture of the Human Pre-mRNA 3□Processing Complex. Mol. Cell 2009, 33 (3), 365–376.

(51) Słabicki, M.; Park, J.; Nowak, R. P.; et al. Expanding the Druggable Zinc-Finger Proteome Defines Properties of Drug-Induced Degradation. Mol. Cell 2025, 85 (16), 3184-3201.e14.

(52) Tian, C.; Sun, L.; Liu, K.; et al. Proteome-Wide Ligandability Maps of Drugs with Diverse Cysteine-Reactive Chemotypes. Nat. Commun. 2025, 16 (1), 4863.

(53) Biggs, G. S.; Cawood, E. E.; Vuorinen, A.; et al. Robust Proteome Profiling of Cysteine-Reactive Fragments Using Label-Free Chemoproteomics. Nat. Commun. 2025, 16 (1), 73.

(54) Wang, Y.; Wang, C. Quantitative Reactive Cysteinome Profiling Reveals a Functional Link between Ferroptosis and Proteasome-Mediated Degradation. Cell Death Differ. 2023, 30 (1), 125–136.

(55) Hughes, C. S.; Moggridge, S.; Müller, T.; et al. Single-Pot, Solid-Phase-Enhanced Sample Preparation for Proteomics Experiments. Nat. Protoc. 2019, 14 (1), 68–85.

(56) Navarrete-Perea, J.; Yu, Q.; Gygi, S. P.; et al. Streamlined Tandem Mass Tag (SL-TMT) Protocol: An Efficient Strategy for Quantitative (Phospho) Proteome Profiling Using Tandem Mass Tag-Synchronous Precursor Selection-MS3. J. Proteome Res. 2018, 17 (6), 2226–2236.

(57) Eng, J. K.; Jahan, T. A.; Hoopmann, M. R. Comet: An Open-Source MS/MS Sequence Database Search Tool. PROTEOMICS 2013, 13 (1), 22–24.

(58) Elias, J. E.; Gygi, S. P. Target-Decoy Search Strategy for Increased Confidence in Large-Scale Protein Identifications by Mass Spectrometry. Nat. Methods 2007, 4 (3), 207–214.

(59) Huttlin, E. L.; Jedrychowski, M. P.; Elias, J. E.; et al. A Tissue-Specific Atlas of Mouse Protein Phosphorylation and Expression. Cell 2010, 143 (7), 1174–1189.

(60) Savitski, M. M.; Wilhelm, M.; Hahne, H.; et al. A Scalable Approach for Protein False Discovery Rate Estimation in Large Proteomic Data Sets [S]. Mol. Cell. Proteomics 2015, 14 (9), 2394–2404.

(61) Gassaway, B. M.; Li, J.; Rad, R.; et al. A Multi-Purpose, Regenerable, Proteome-Scale, Human Phosphoserine Resource for Phosphoproteomics. Nat. Methods 2022, 19 (11), 1371–1375.

(62) Beausoleil, S. A.; Villén, J.; Gerber, S. A.; et al. A Probability-Based Approach for High-Throughput Protein Phosphorylation Analysis and Site Localization. Nat. Biotechnol. 2006, 24 (10), 1285–1292.

(63) Xu, S.; Hu, E.; Cai, Y.; et al. Using clusterProfiler to Characterize Multiomics Data. Nat. Protoc. 2024, 19 (11), 3292–3320.

(64) Shannon, P.; Markiel, A.; Ozier, O.; et al. Cytoscape: A Software Environment for Integrated Models of Biomolecular Interaction Networks. Genome Res. 2003, 13 (11), 2498–2504.

