## Supplemental Figure S1-6 for "Analysis of Confounding Factors in Reactive Cysteine Profiling Reveals Enhanced Chromatin-Protein Association via CDK7 Inhibition by THZ1"

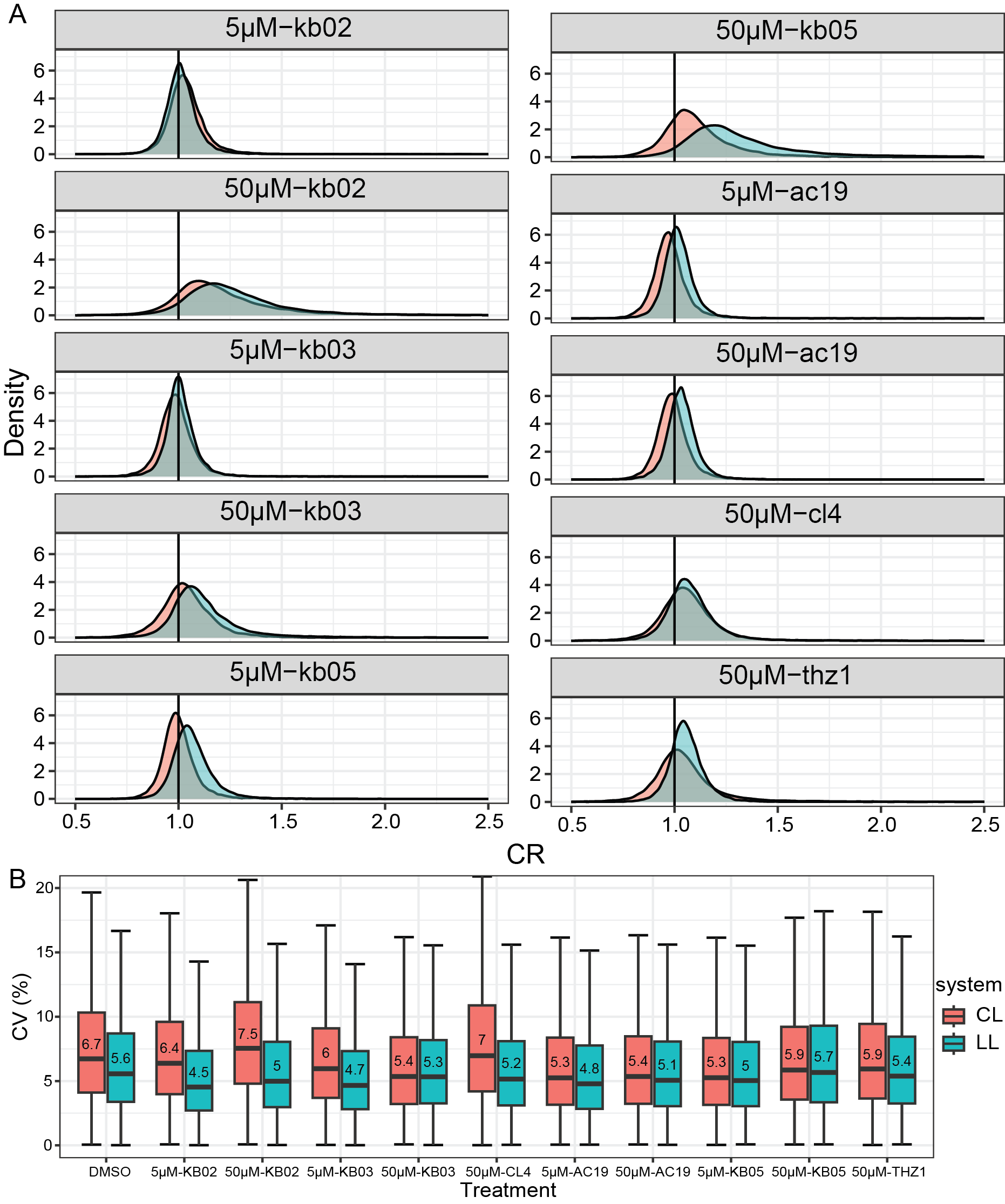

**Figure S1. A)** Competition ratio distribution across two systems under different drug treatments. **B)** Coefficient of variation (CV) of cysteine measurements. Numbers represent median values.

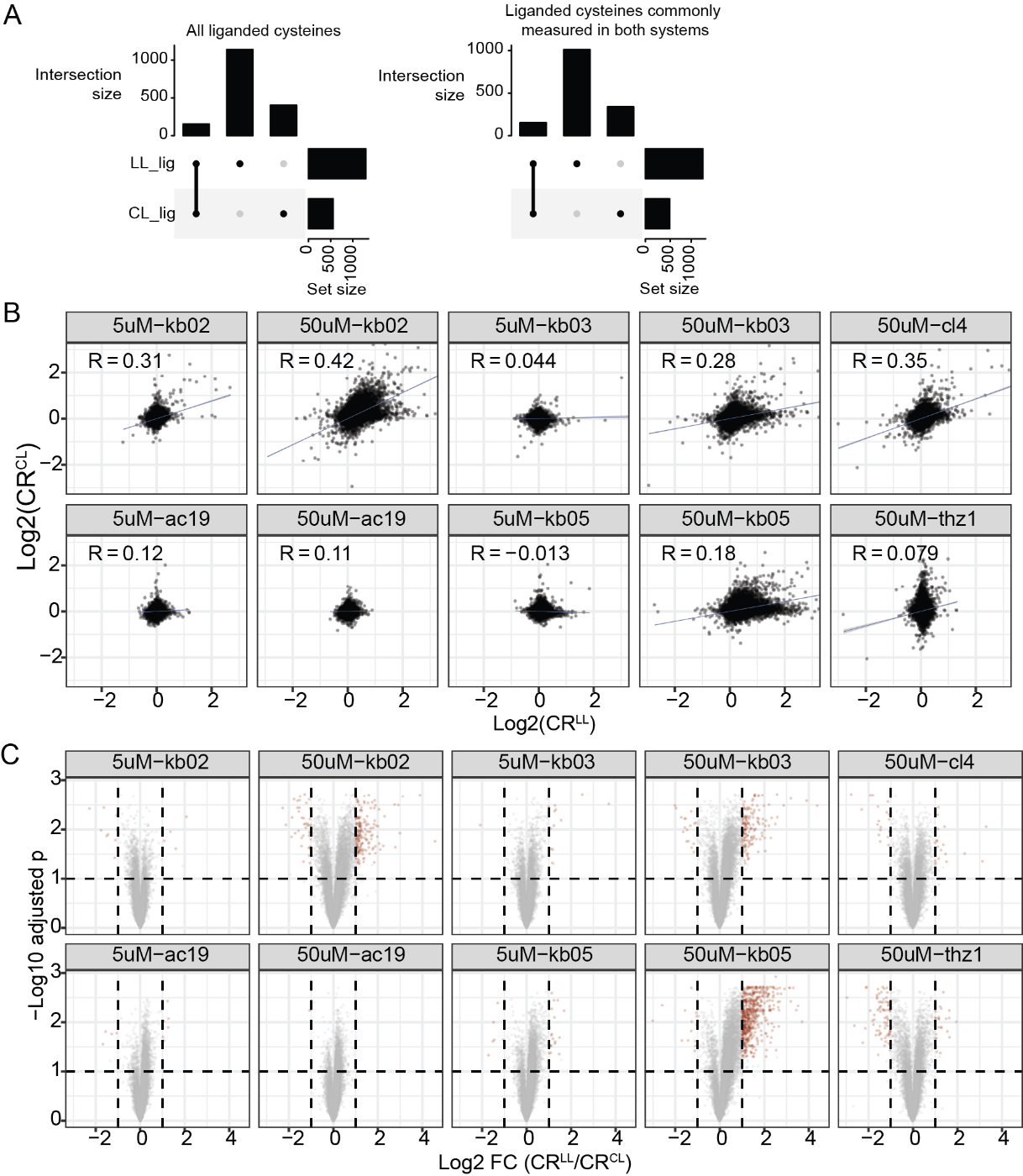

**Figure S2. Comparison of ligandable cysteines across two systems. A)** Overlap of liganded cysteines. All liganded cysteines are included in the left panel and liganded cysteines commonly measured by both systems are in the right panel. **B)** Correlation between CR^CL^ and CR^LL^ by each treatment. **C)** Volcano plot of fold changes between CR^CL^ and CR^LL^. Vertical dotted lines represent 2 fold changes, and horizontal dotted lines represent adjusted p=0.1.

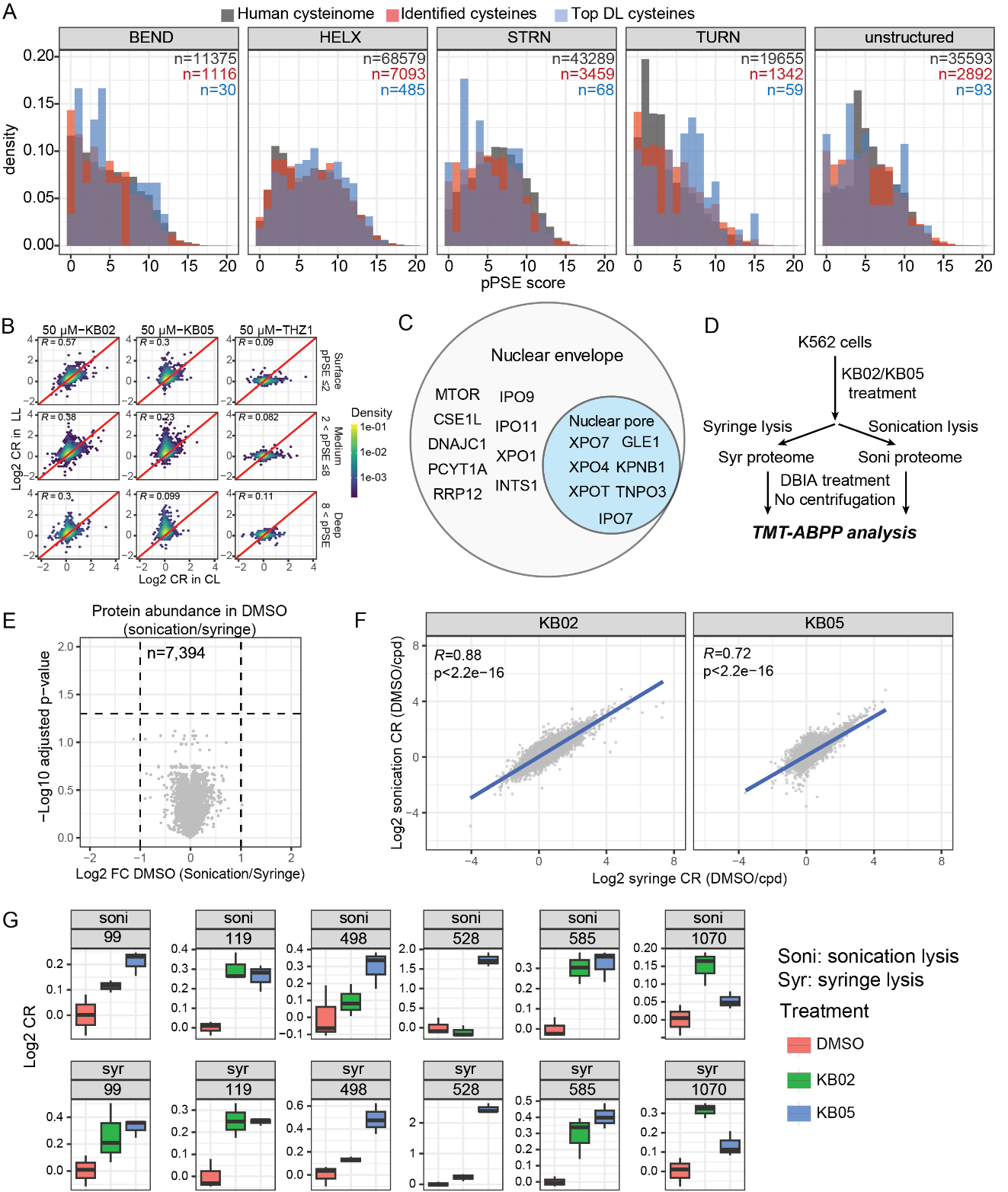

**Figure S3. A)** pPSE distribution of all human cysteines with confident AlphaFold2 prediction (pLDDT>70), quantified cysteines in this study, and cysteines with different ligandability profiles between CL and LL, stratified by substructure types. **B)** Correlation of CR^LL^ vs. CR^CL^ of cysteines of three categories determined by pPSE score. **C)** Proteins containing DL cysteines were identified in GO terms shown in Fig 3E. **D)** Workflow to compare the post-treatment lysis methods in CL experiments. **E)** Proteome abundance between DMSO samples using syringe vs. sonication lysis. **F)** Correlation (Person r) of CR values using syringe vs. sonication lysis. **G)** Ligandability profiles of all measured XPO1 cysteines.

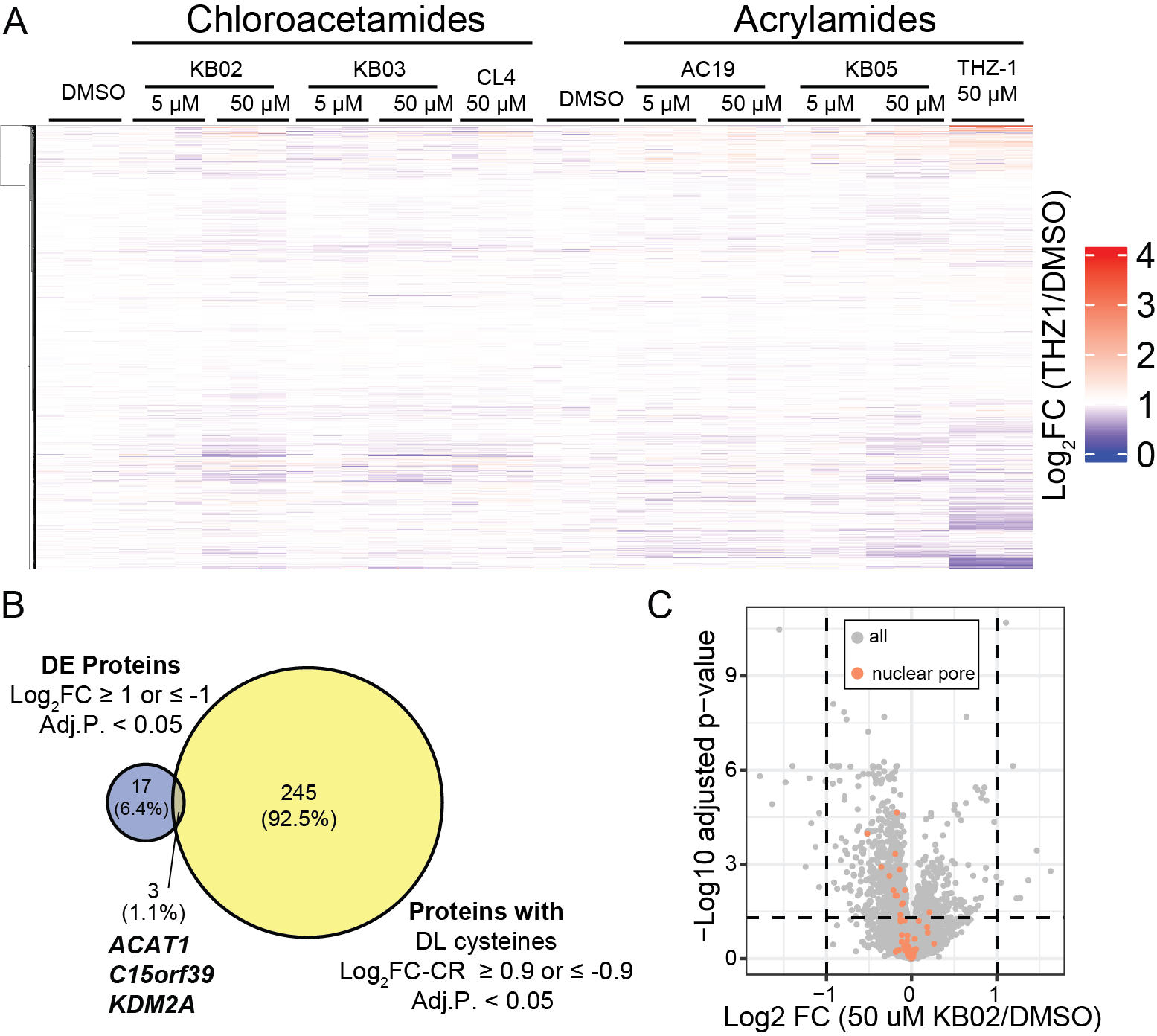

**Figure S4. Protein abundance changes induced by compound treatment in live cells. A)** Protein abundance change relative to DMSO measured after drug treatment in each live-cell ABPP sample. **B)** Overlap of DE proteins and Proteins containing DL cysteines by KB02. **C)** Volcano plot comparing 50 uM KB02 to DMSO. Nuclear pore proteins are colored orange.

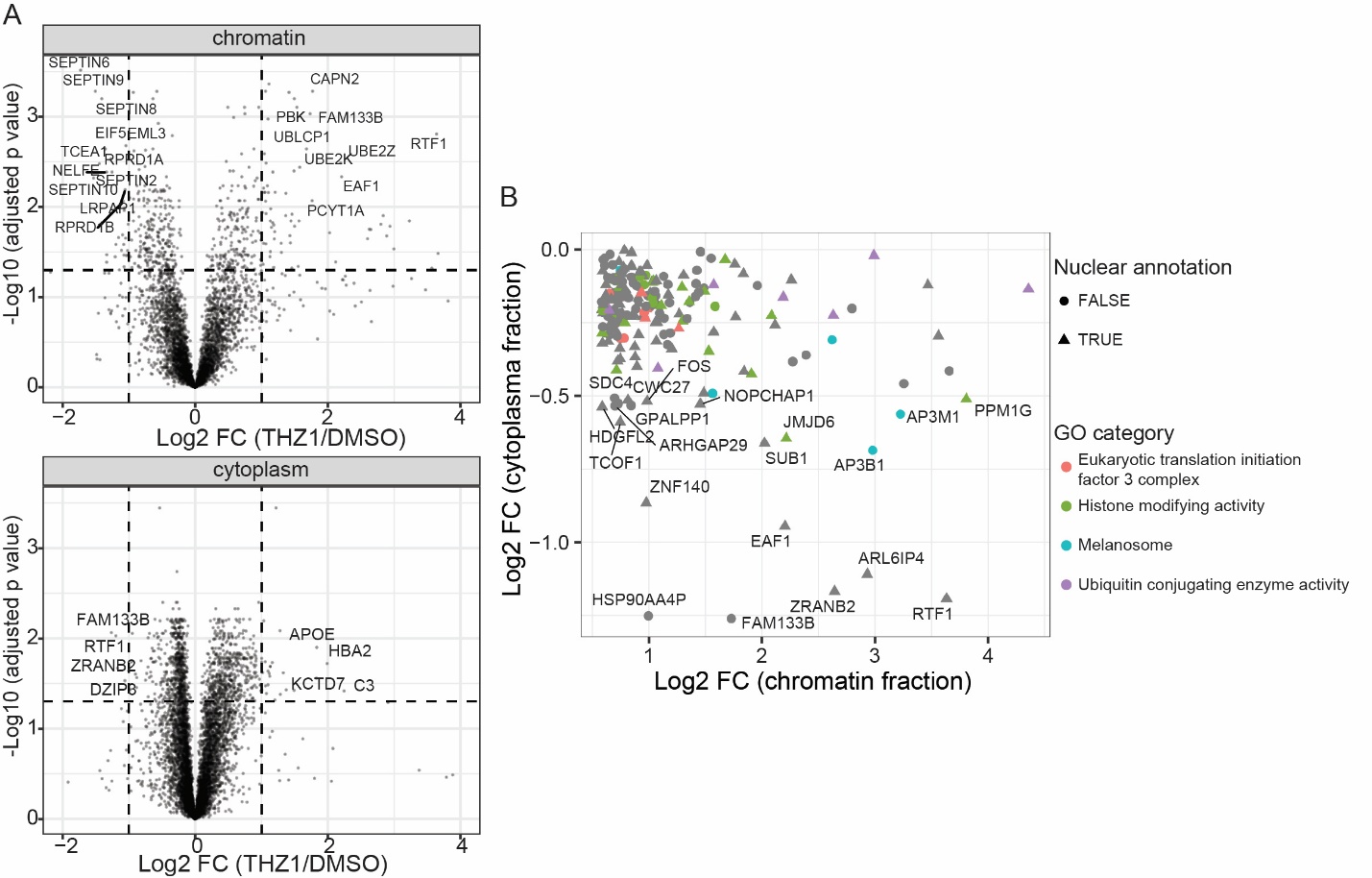

**Figure S5. Subcellular proteomics with THZ1.** **A)** volcano plots of protein changes in chromatin and cytoplasm fractions. **B)** all proteins with reduced abundance in cytoplasm and increased abundance in chromatin. Proteins with annotated nuclear location are triangles.

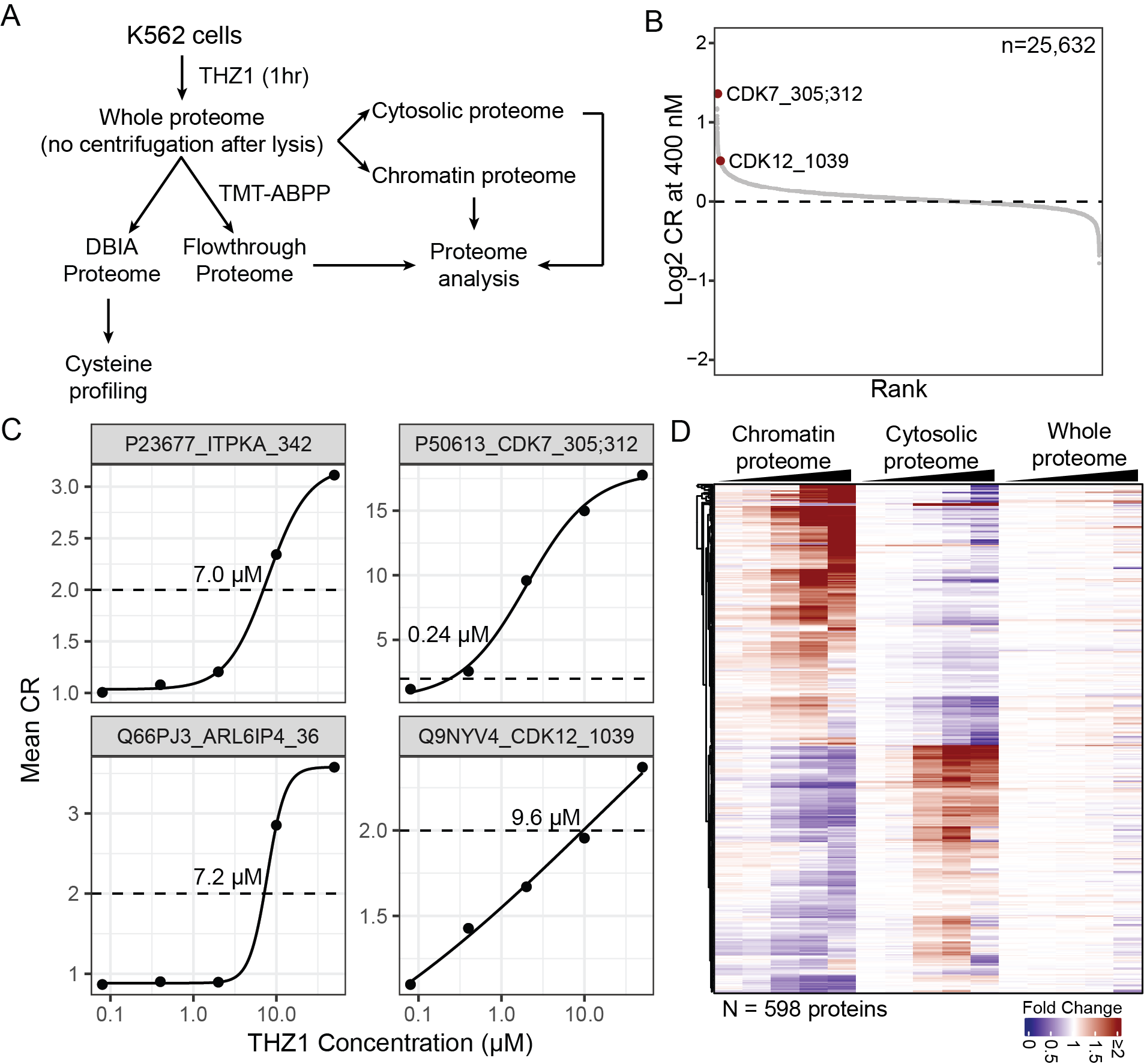

**Figure S6. Subcellular proteomics across a THZ1 dose range. A)** Workflow for cysteine and proteome profiling of K562 cells treated with THZ1 at concentrations ranging from 80 nM to 50 µM for 1 hour. **B)** Ranked CR values of cysteines at 400 nM THZ1. The quantified CDK7 peptide bears two DBIA-labeled cysteines, C305 and C312, where C312 is the THZ1 target site, and is therefore annotated as CDK7_305;312 on the plot. **C)** Dose-response curves of on- and off-target cysteines. Annotated concentrations indicate the concentration at which 50% cysteine occupancy is achieved. **D)** Heatmap of significantly changed proteins (adjust ANOVA P < 0.001) under THZ1 treatment in at least one of three proteomic datasets: chromatin, cytosolic, or whole proteome.
